# Thyroid Hormone Receptor Beta Signaling is a Targetable Driver of Prostate Cancer Growth

**DOI:** 10.1101/2025.03.05.641137

**Authors:** Aleksandra Fesiuk, Daniel Pölöske, Elvin D. de Araujo, Geordon A. Frere, Timothy B. Wright, Gary Tin, Ji Sung Park, Yasir S. Raouf, Olasunkanmi O. Olaoye, Michaela Schlederer, Sandra Högler, Cécile Philippe, Heidi A. Neubauer, Osman Aksoy, Adam Varady, Alejandro Medaglia Mata, Maxim Varenicja, Boglárka Szabó, Theresa Weiss, Gabriel Wasinger, Torben Redmer, Martin Susani, Clemens P. Spielvogel, Jing Ning, Maik Dahlhoff, Martin Schepelman, Richard Kennedy, Geoffrey Brown, Jenny Persson, Vojtech Bystry, Oldamur Hollóczki, David M. Heery, Patrick T. Gunning, Olaf Merkel, Brigitte Hantusch, Lukas Kenner

## Abstract

Thyroid hormone (TH) signaling plays a major role in the development, energy homeostasis, and metabolism of most tissues. Recent observations have identified THs as drivers of prostate cancer (PCa) tumor development and progression. We reported that the T3-scavenger protein µ-crystallin (CRYM) regulates the development and progression of PCa and that this involved crosstalk with the androgen receptor (AR) signaling. However, the mechanisms remain incompletely understood. Here, we explored the role of thyroid hormone receptor β (TRβ), which is the main effector of TH signaling, in the context of PCa. The use of the TRβ-selective antagonist NH-3 inhibited PCa cell proliferation *in vitro* and reduced tumor size in PCa xenograft models. Notably, NH-3 was highly effective in the engrafted 22Rv1 cell line, a model for castration-resistant PCa (CRPC). Mechanistic studies revealed that NH-3 downregulates AR and the AR target genes *Nkx3.1* and *KLK3* (*PSA*). NH-3 was a more effective anticancer agent than enzalutamide and showed synergistic properties in combined use. Evidence from human datasets corroborates our findings whereby elevated TRβ expression and mutations in TH signaling pathways are associated with the onset of PCa. Collectively, these results establish TRβ as a mediator of tumorigenesis in PCa and identify NH-3 as a promising therapeutic agent for targeting AR signaling, particularly in CRPC.

## INTRODUCTION

Prostate cancer (PCa) is the second most frequent malignancy in men (Jemal et al., 2010) and the sixth leading cause of death worldwide (Sung et al., 2021). Epidemiological studies estimate that one in eight men will be diagnosed with PCa during their lifetime (Jain et al., 2025). Disease progression highly depends on androgen receptor (AR) signaling, which regulates cell proliferation and survival. The AR, a nuclear receptor and ligand-dependent transcription factor, is activated by testosterone and dihydrotestosterone (DHT). Given its central role in PCa growth, most therapeutic strategies target AR signaling. The first-line treatment of PCa is androgen deprivation therapy (ADT), which suppresses testosterone and DHT either through surgical castration or anti-androgen drugs (Wade and Kyprianou, 2018). Second-generation AR inhibitors, such as enzalutamide, further block AR activity (Culig, 2017). However, up to 20% of patients develop resistance to ADT and progress to castration-resistant prostate cancer (CRPC) (Vellky and Ricke, 2020). Nearly 50% of CRPC cases advance to metastatic CRPC (mCRPC) within three years of diagnosis (Akaza et al., 2018), resulting in a poor prognosis and a diminished quality of life.

Recent studies have shown that THs regulate the pituitary-gonadal axis and androgen signaling (Anguiano et al., 2022) and linked altered serum levels of THs and thyroid-stimulating hormone (TSH) with the risk of PCa (Şenel et al., 2020; Mondul et al., 2012; Ovčariček et al., 2020). At a cellular level, THs promote PCa cell proliferation *in vitro* (Anguiano et al., 2022; Tsui et al., 2008; Zhang et al., 1999) and tumor progression *in vivo* ( Miro et al., 2022).

TH signaling has been implicated in PCa development and progression by stimulating cell proliferation and tumor growth. However, the molecular mechanisms underlying this process remain unclear.

The recent FDA approval of the TRβ-selective agonist resmetirom for treating non-alcoholic steatohepatitis (NASH) highlights that the therapeutic potential of targeting TRβ (Harrison et al., 2024; Keam, 2024) is recognized.

Our previous findings identified reduced expression of the T3-scavenger protein µ-crystallin (CRYM) in aggressive PCa, correlating with earlier biochemical recurrence and poor survival (Aksoy et al., 2021). CRYM is a negative regulator of T3 availability, implying enhanced T3-mediated signaling when CRYM is downregulated (Aksoy et al., 2022). CRYM overexpression reduces TRβ expression, suggesting regulatory crosstalk with AR signaling, evidenced by suppressed PSA (prostate-specific antigen, KLK3). Based on these findings, we hypothesize that TRβ is a key driver of PCa progression and a potential therapeutic target.

In this study, we investigated the role of TRβ in PCa proliferation and tumor development. We demonstrate that the TRβ-selective antagonist NH-3 significantly reduced PCa cell proliferation in vitro and suppressed tumor growth in xenograft models *in vivo*. NH-3 treatment led to AR degradation and loss of AR-dependent genes. Analysis of human datasets revealed enhanced TRβ expression on mRNA and protein levels in PCa compared to normal tissue, providing further evidence for a tumor-driving role of TRβ in PCa patients. Our results show that TRβ blockade is a promising therapeutic approach, particularly for patients with ADT resistance.

## RESULTS

### Thyroid hormone signaling drives PCa cell proliferation

To investigate the role of TRβ in PCa, we analyzed TRβ protein expression within a panel of PCa cell lines. RWPE1 cells, derived from histologically normal prostate tissue, served as a non-malignant model, while BPH-1 cells, derived from benign prostatic hyperplasia (BPH), represented premalignant prostate tissue. LNCaP and 22Rv1 served as AR-positive models. Consistent with prior studies (Tindall and Lonergan, 2011), only these two cell lines expressed the full-length AR, with the AR-V7 form present in 22Rv1 (**Fig. 1A**, middle panel), which makes them a model for castration-resistant prostate cancer (CRPC). PC3 and DU-145 cells, which are AR-negative, were utilized as aggressive PCa cell lines. TRβ protein expression was detected in all cell lines analyzed (**Fig. 1A**, upper panel). Given their AR expression profiles, LNCaP and 22Rv1 were selected for subsequent experiments, with 22Rv1 chosen specifically for their relevance as a CRPC model.

**Figure 1.**
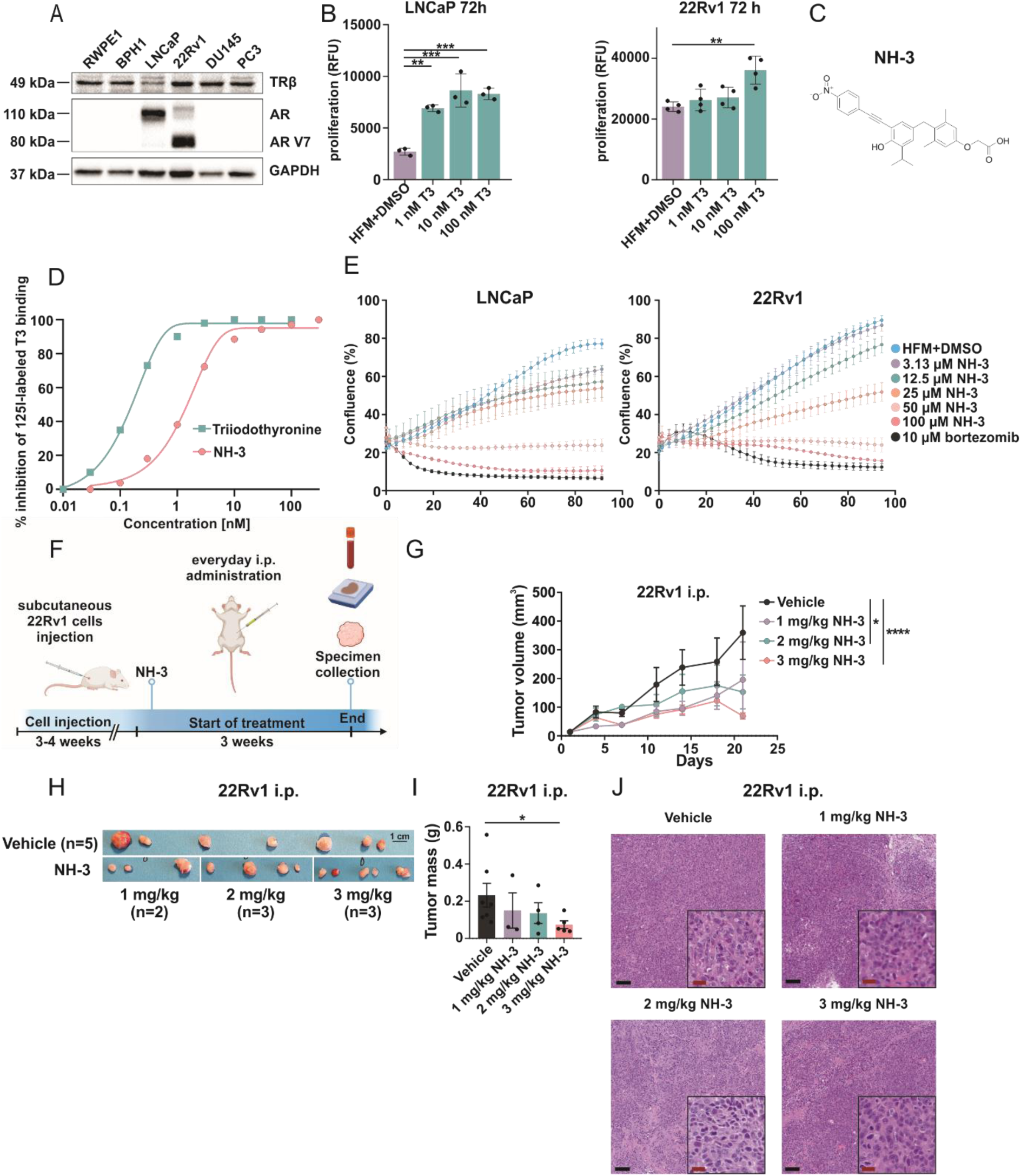
Treatment of PCa cells with TRβ-specific inhibitor NH-3 reduces PCa cell proliferation *in vitro* and xenograft tumor growth *in vivo*. **(A)** TRβ and AR protein expression in PCa cell lines RWPE1, BPH-1, LNCaP, 22Rv1, PC3 and DU145. GAPDH as protein loading control, AR V7 – AR splice variant 7. **(B)** T3 stimulates LNCaP and 22Rv1 proliferation in hormone-free conditions in a dose-dependent manner. **(C)** Chemical structure of NH-3. **(D)** Inhibition of the binding of [I^125^]-radiolabeled T3 to TRβ by unlabeled T3 and NH-3. Ki of T3 = 0.15 nM, Ki of NH-3 = 1.31 nM. **(E)** Reduced proliferation of LNCaP and 22Rv1 as measured every 2 h upon NH-3 treatment in hormone-free conditions as calculated by % cell confluence in each well. Bortezomib as positive control for cell death. **(F)** Scheme of NH-3 i.p. treatment in the 22Rv1 xenograft model. Created with BioRender.com. **(G)** Tumor volume during the treatment. **(H)** Visual representation of 22Rv1 xenografts treated i.p. with 1, 2, 3 mg/kg/day of NH-3 via i.p. administration. **(I)** Tumor mass after the treatment. **(J)** HE staining of tumors. Statistical analysis with 2-way ANOVA with multiple comparisons. Mean ± SD, *p < 0.05, **p < 0.01, and ***p < 0.001.

To determine whether TRβ signaling influenced proliferation, we treated LNCaP and 22Rv1 cells with T3 under hormone-free medium (HFM) conditions. In agreement with previous reports, T3 stimulation significantly increased the proliferation of LNCaP cells (Esquenet et al. 1995; Tsui et al., 2008; Zhang et al., 1999). Notably, we extended these findings by demonstrating that 22Rv1 cells also exhibited increased proliferation at 72 h of T3 treatment (**Fig.1B**). These results further support the role of thyroid hormone signaling in driving PCa cell proliferation, including CRPC models.

### NH-3 is a selective ligand for TRβ

To investigate the role of TRβ in PCa cell proliferation, we utilized the synthetic TRβ-selective ligand NH-3 **(Fig. 1C**, chemical structure) (Nguyen et al., 2002). NH-3, a potent and specific TRβ antagonist, was used in both cell culture and various animal models (Lim et al., 2002; Walter et al., 2021; Grover et al., 2007). We first reexamined the specificity of NH-3 for TRβ antagonism to exclude possible additional interactions and off-target effects. Binding assays of TRβ, the TRβ binding partners retinoid X receptor Alpha (RXRα), retinoid X receptor Beta (RXRβ), and four hormone receptors androgen receptor (AR), estrogen receptor (ER), glucocorticoid receptor (GR), progesterone receptor (PR) that are activated by cholesterol-derived hormones and have known roles in PCa (Shiota et al., 2019) were chosen. In each assay, the binding of respective radioactively labeled ligands to its receptor was monitored in the presence of increasing NH-3 concentrations. NH-3 only led to inhibition of T3 binding to TRβ (IC50 = 1.68 nM), with no inhibition of any other ligand binding to respective receptors (**Fig.1D**).

### NH-3 reduced the proliferation of PCa cells *in vitro*

To assess in detail the impact of TRβ inhibition on PCa cell viability, IC_50_ values of NH-3 for the human PCa cell lines were determined (**Suppl. Fig. 1A**). Cells were treated with increasing concentrations of NH-3 in hormone-free (HFM) conditions, and cell growth was monitored every two hours using live cell microscopy. NH-3 treatment significantly reduced the cell layer coverage of LNCaP and 22Rv1 cells as compared to the vehicle control (**Fig. 1E**). Further analysis using flow cytometry to assess the proliferation marker Ki-67 revealed a shift from Ki-67^high^ cells towards a Ki-67^low^ population, indicating suppressed proliferation rates **(Suppl. Fig. 1B)**. To confirm these findings, a resazurin assay was used as an independent cell viability readout based on metabolic activity. NH-3 treatment led to the inhibition of the viability of LNCaP and 22Rv1 cells in a concentration-dependent manner (**Suppl. Fig. 1C**, upper panel). Importantly, to rule out the possibility that the observed growth inhibition resulted from TRβ ligand deprivation, NH-3 was also tested under full-media (FM) conditions (**Suppl. Fig. 1C**, bottom panel). Both cell lines exhibited similar antiproliferative effects, proving NH-3 efficacy in physiological conditions.

These findings establish TRβ blockade by NH-3 as a potent strategy to inhibit PCa cell proliferation, reinforcing the pivotal role of TRβ in modulating cellular growth processes in PCa.

### NH-3 treatment reduced 22Rv1 PCa cell xenograft growth

We extended our investigation of NH-3 function to PCa xenograft models *in vivo*. Given the strong effect observed in the 22Rv1 cell line and its relevance as a CRPC model, we first used this cell line for xenograft experiments. NOD scid gamma (NSG) mice were subcutaneously injected with 22Rv1 cells, and treatment started once the tumors had reached approximately 100 cm^3^. For three weeks, mice received a daily intraperitoneal (i.p.) injection of NH-3 at doses of 1, 2, or 3 mg/kg. Tumor size and body weight were monitored throughout the treatment period, and tumors, sera, and tissue samples were collected at study termination (**Fig. 1F**). NH-3 treatment led to dose-dependent tumor growth inhibition (TGI), with 80% tumor growth inhibition (TGI) at 3 mg/kg, 57% TGI at 2 mg/kg, and 46% TGI at 1 mg/kg. Notably, tumors in the 3 mg/kg group exhibited sustained suppression, with volumes remaining stable at approximately 125 mm³ through the study endpoint (**Fig. 1G, H, I, J**). NH-3 treatment was well tolerated, as indicated by the absence of any significant body weight changes (**Suppl. Fig. 1D**). Serum analysis revealed no abnormalities in metabolic markers, ruling out potential liver or kidney toxicity (**Suppl. Fig. 1E**). Histopathological examination of heart, lung, kidney, liver, spleen, and colon tissues showed no morphological abnormalities in the 3 mg/kg group compared to vehicle. These findings highlight NH-3 as a potent and well-tolerated TRβ antagonist that effectively suppresses CRPC tumor growth *in vivo*.

NH-3 has been reported to be bioavailable for oral administration (Nguyen et al., 2002). Therefore, we tested its antitumor efficacy when administered by oral gavage. After establishing 22Rv1 xenograft tumors, mice received daily oral gavage NH-3 for two weeks at 3 mg/kg and 6 mg/kg, along with a vehicle control group (**Fig. 2A**). Tumor size and body weight were monitored as previously described. At the end of the treatment, NH-3-treated mice exhibited significantly smaller 22Rv1 tumors with reduced mass and volume compared to controls (**Fig. 2B, C, D, E**). Both treatment concentrations resulted in 50-55% TGI, while 6 mg/kg did not show higher efficacy than 3 mg/kg, indicating a minimal effective dose beyond which no further enhancement of tumor reduction occurs. Importantly, mice maintained stable body weight throughout the 2-week treatment period (**Suppl. Fig. 2A**). Serum analyses confirmed no signs of kidney or liver dysfunction (**Suppl. Fig. 2B**), and histopathological evaluation showed no degenerative or inflammatory lesions in heart, lung, kidney, spleen, and colon of treated animals compared to vehicle, supporting the safety profile of NH-3. Taken together, the above findings have shown that NH-3 inhibits PCa xenograft growth when administered either orally or i.p. and that continual dosing was well-tolerated with no detectable toxicity.

**Figure 2.**
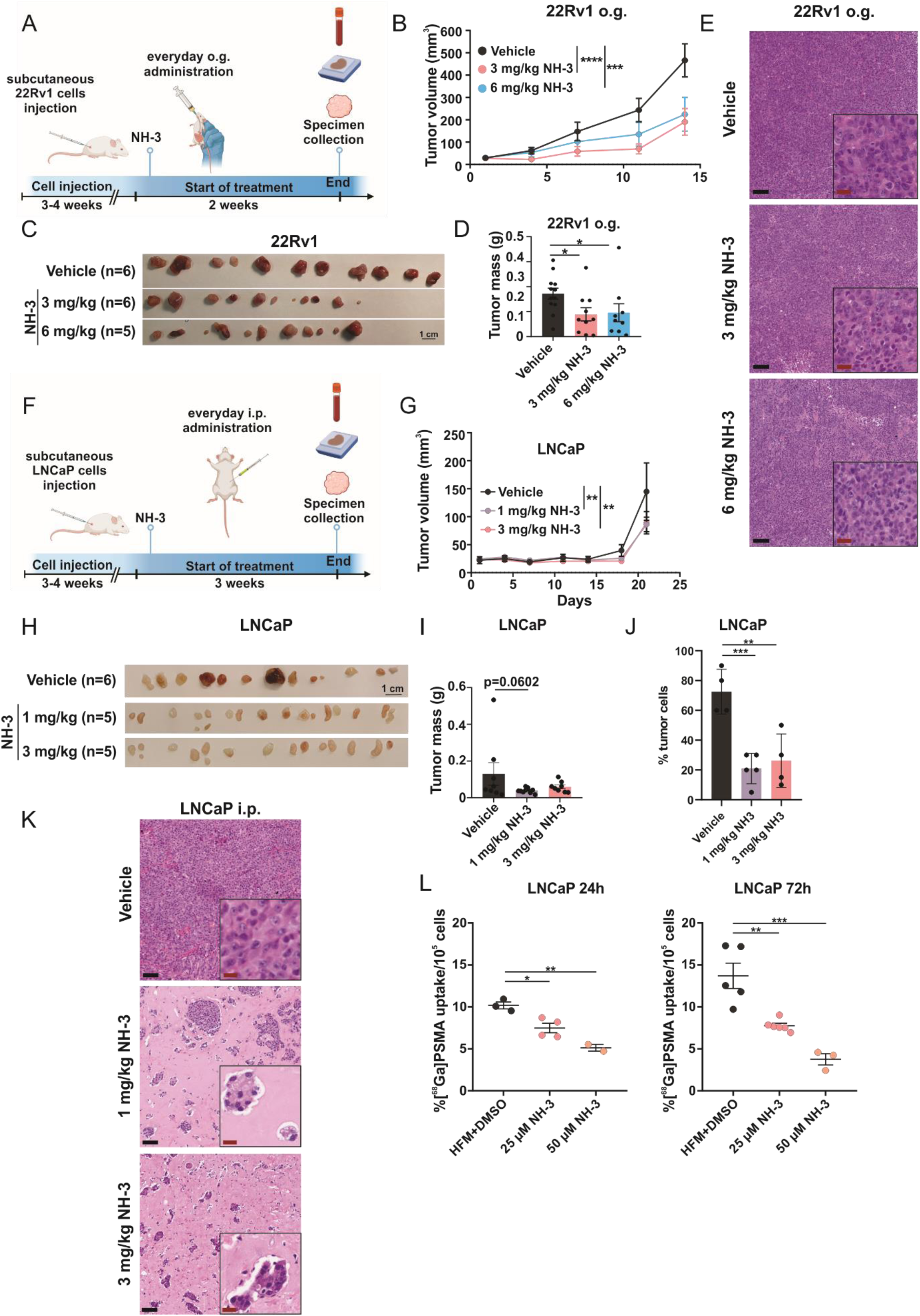
Inhibition of LNCaP and 22Rv1 PCa tumor growth by TRβ-specific inhibitor NH-3 *in vivo* and by different administration routes. **(A)** Scheme of NH-3 oral gavage (o.g.) treatment in 22Rv1 xenograft model. Created with BioRender.com. **(B)** Reduced tumor volume during treatment. **(C)** Visual representation of 22Rv1 xenografts in mice treated with 3 and 6 mg NH-3/kg/day by o.g. **(D)** Reduced tumor mass after the treatment endpoint. **(E)** H&E staining of tumors. **(F)** Scheme of NH-3 intraperitoneal (i.p.) treatment in LNCaP xenograft mice model. Created with BioRender.com. **(G)** Reduced tumor volume during the treatment. **(H)** Visual representation of LNCaP xenografts treated with 1 and 3 mg NH-3/kg/day via i.p. administration. **(I)** Reduced tumor mass after the treatment endpoint. **(J)** Percentage of tumor cells in LNCaP tumors treated with vehicle control, 1 and 3 mg/kg/day NH-3. **(K)** H&E staining of tumors. **(L) G**amma counter measurements of [^68^Ga]PSMA uptake in LNCaP cells treated with vehicle control, 25 µM and 50 µM NH-3. Results shown as % uptake and normalized to 10^5^ cells. Mean ± SD, *p < 0.05, **p < 0.01, and ***p < 0.001.

### NH-3 treatment reduced LNCaP PCa xenograft growth and PSMA uptake

The findings for the 22Rv1 xenograft model were validated using LNCaP xenografts. Following cell engraftment and over a three-week period, mice received daily i.p. injections of NH-3 at 1 mg/kg and 3 mg/kg (**Fig. 2F**). Tumors in the NH-3 treated group were significantly smaller I volume (**Fig. 2G, H**), and a trend in reduction of tumor mass was observed (**Fig. 2I**). They were less palpable and barely visible through the skin, which limited precise monitoring of growth progression. Treatment resulted in 40% TGI in both tested NH-3 concentrations, indicating a slightly smaller effect compared to that seen for 22Rv1 xenografts. The pathologist’s histological evaluation based on H&E staining revealed reduced tumor cell percentage in NH-3-treated mice relative to vehicle controls (**Fig.2J, K**). NH-3 treated mice did not lose body weight throughout the experiment (**Suppl. Fig. 2C**), serum analysis showed no signs of toxicity (**Suppl. Fig. 2D**), and histopathological analysis did not reveal any signs of tissue degeneration or inflammation.

Prostate-specific membrane antigen (PSMA) is a transmembrane glycoprotein highly expressed in PCa. It is widely used as a diagnostic marker for PCa, and its elevated expression is indicative of disease progression (Tindall and Lonergan, 2011). LNCaP cell line expresses high levels of PSMA (Fan et al., 2015; Gorges et al., 2016) and serves as a valuable model for PSMA research in PCa. Using a Gamma counter, we quantified the *in vitro* uptake of [^68^Ga]PSMA radiotracer in NH-3-treated LNCaP cells. A dose-dependent reduction of radiotracer uptake was observed at 25 µM and 50 µM NH-3 concentrations after 24 h and 72 h of treatment (**Fig.2L**). This effect underscores NH-3’s impact on PSMA-associated signaling pathways and tumor metabolism and further indicates reduced tumor growth. No nonspecific binding was detected, confirming the specificity of [^68^Ga]PSMA under the experimental conditions. Together, these findings demonstrate that NH-3 inhibits PCa xenograft growth through both oral and i.p. administration, with well-tolerated dosing and no detectable toxicity. Additionally, NH-3 reduces PSMA expression and activity.

### TRβ inhibition affects the AR and AR-regulated gene expression

AR-signaling is the pivotal pathway driving PCa progression and the transition to CRPC disease (Tindall and Lonergan, 2011). T3 was observed to stabilize AR levels in LNCaP cells (Esquenet et al., 1995). We examined the influence of T3 stimulation on TRβ and AR expression in LNCaP cells under hormone-free medium (HFM) conditions. While TRβ protein levels remained stable, HFM led to reduced AR expression, which was fully restored upon T3 treatment at later time points (**Fig. 3A**).

**Figure 3.**
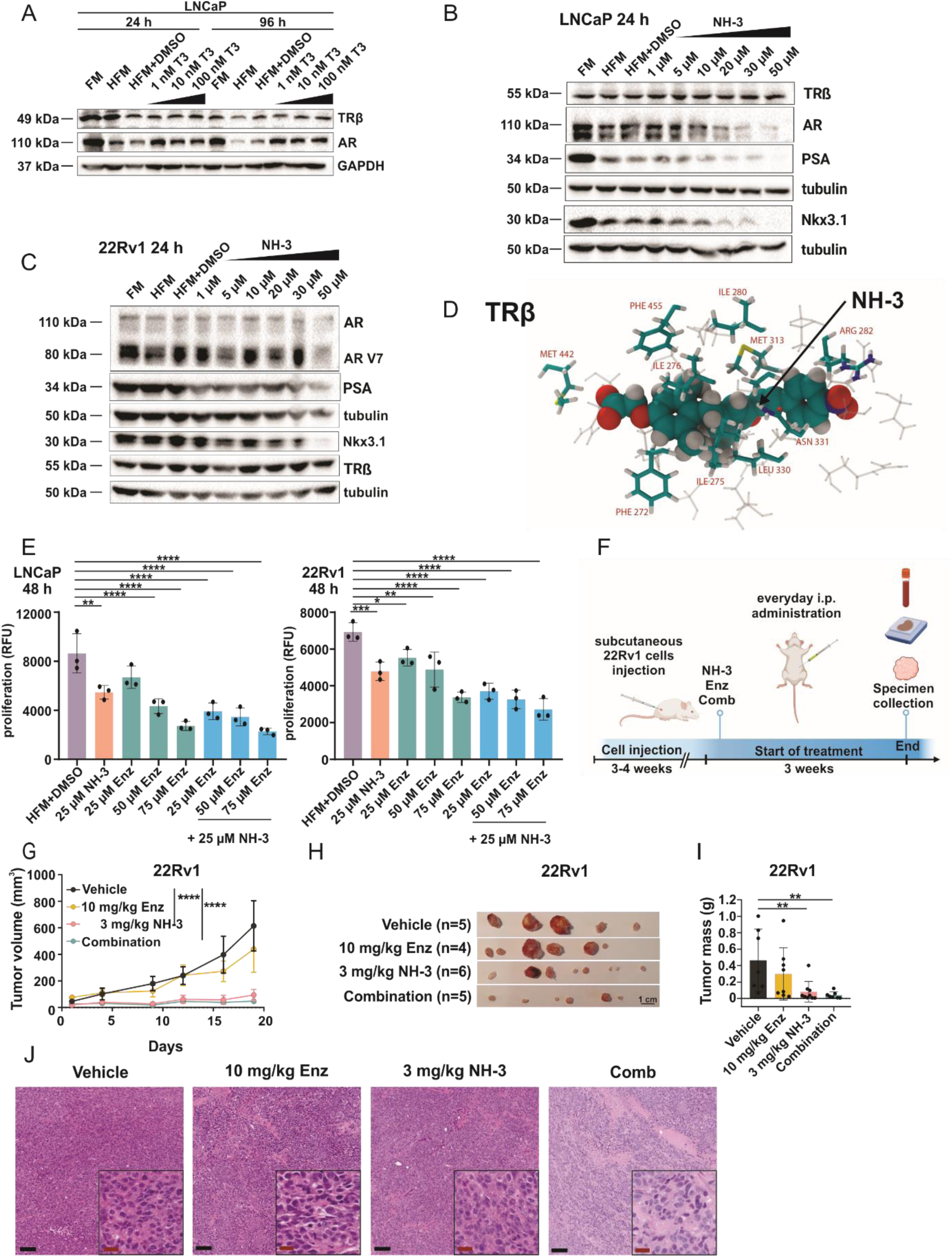
Selective TRβ blockade induces downregulation of AR and AR-regulated genes and is more effective than enzalutamide. **(A)** Western blot analysis of LNCaP cells treated with 1 nM, 10 nM, 100 nM T3 after 24 and 96 h. Representative protein expression of TRβ, AR, GAPDH as control; FM – full media, HFM – hormone-free media. **(B)** Western blot analysis of LNCaP cells treated with increasing NH-3 concentrations after 24 h. Representative blot showing expression of TRβ, AR, PSA, Nkx3.1; GAPDH and tubulin as loading controls. **(C)** Western blot analysis of 22Rv1 cells treated with increasing NH-3 concentrations after 24. Representative protein expression of TRβ, AR, PSA, Nkx3.1; GAPDH and tubulin as loading controls. **(D)** Binding of NH-3 to the ARß (2PIN), as obtained from molecular docking calculations and subsequent molecular dynamics simulations. The residues of the protein that are adjacent to the ligand are highlighted and labelled. **(E)** Proliferation (RFU) of LNCaP and 22Rv1 cells treated with 25 µM NH-3, 75, 57, 25 µM enzalutamide and combination treatment; Enz – enzalutamide. **(F)** Scheme of NH-3, enzalutamide and combination i.p. treatment in 22Rv1 xenograft model. Created with BioRender.com. **(G)** Reduced tumor volume during the treatment. **(H)** Visual representation of 22Rv1 xenografts treated i.p. with 3 mg/kg/day NH-3, 10 mg/kg/day enzalutamide and combinatory treatment endpoint. **(I)** Reduced tumor mass after the treatment. **(J)** H&E staining of tumors.

We evaluated whether NH-3 treatment modulates TRβ, AR and its downstream target gene expression. TRβ protein levels remained stable in LNCaP cells with increasing concentrations of NH-3 (**Fig. 3B, C**). These findings confirmed that NH-3 operates via inhibition of TRβ activity rather than a ligand-induced degradation mechanism observed for other nuclear receptor modulators (Salami et al., 2018; Gustafson et al., 2015; Chen et al., 2024). Notably, NH-3 treatment led to reduced AR protein levels after 24 h (**Fig. 3B, C**), which was sustained at 72 h (**Suppl. Fig. 3 A, B**). In 22Rv1 cell line also AR-V7 downregulation was observed. Additionally, the AR downstream targets Nkx3.1 and PSA were downregulated in in a dose-dependent manner (**Fig. 3B, C**). These findings showed that TRβ antagonism by NH-3 suppresses AR signaling.

To observe how the NH-3 binds to TRβ, we performed molecular docking calculations, supplemented with a sequence of molecular dynamics (MD) simulations. We chose two protein structures from literature, 3GWS (Nascimento et al., 2006) and 2PIN (Estébanez-Perpiñá et al., 2007), both obtained by X-ray diffraction. Each of these structures contains a ligand, the former triiodothyronine (T3), the latter the inhibitor metabolite tiratricol (4HY), clearly defining the native binding site of this receptor. After an initial 10 ns MD run to remove all energetic hotspots from the system, the T3 and 4HY were cut out of the proteins, and the NH-3 molecule was docked into the binding site. Reassuringly, the docking resulted in an identical orientation of the drug inside the protein in both cases. The structure is shown in **Fig. 3D**, with the amino acid side chains adjacent to NH-3 highlighted and labelled. The binding energy was found to be significant, ΔE = −8.7 and −7.0 kcal mol-1 for 3GWS and 2PIN, respectively. Performing 20 ns of MD simulations on the system allowed the further adjustment of the conformations of the biomolecules and NH-3 to ensure an even stronger binding. The strength of the interaction can be represented by the development of interaction energies between the proteins and the drug, averaging at ΔE_int_ = −73.5 and −69.3 kcal mol-1 through the last 10 ns of the simulations for 3GWS and 2PIN, respectively. Cutting out the drug from the resulting structures and performing the docking calculations anew resulted in even more pronounced binding energies, amounting to ΔE = −11.3 and −11.5 kcal mol-1. Whereas these results support the experimental findings regarding a strong interplay between TRβ and NH-3, the fact that the procedure produced quantitatively such similar results for the interaction energies and final, binding energies for two different starting structures indicates the validity of the calculations.

To confirm the observed effects of NH-3 on AR expression, we made use of another TRβ antagonist, 1-850 (**Suppl. Fig. 3C**). 1-850 also inhibited the proliferation of LNCaP and 22Rv1 cells (for IC50 curves see in **Suppl. Fig. 3D**). Treatment of LNCaP cells with 1-850 similarly reduced AR and PSA expression (**Fig. 3E**), emphasizing the role of TRβ role in maintaining AR signaling.

We investigated the therapeutic potential of combining the use of NH-3 with the AR antagonist enzalutamide in LNCaP and 22Rv1 cells. Enzalutamide alone reduced proliferation in a dose-dependent manner, while co-treatment with NH-3 enhanced this effect (**Fig. 3E**). Synergy analysis experiments demonstrated additive growth inhibition across dose-response matrices for both cell lines, as revealed by distinct peaks in 3D synergy maps (**Suppl. Fig. 3F**). Given the synergy observed *in vitro*, we evaluated NH-3 and enzalutamide in vivo using NSG mice engrafted with 22Rv1 cells (**Fig. 3F**). The most effective dose of NH-3 was chosen for combinatory treatments. Mice treated with NH-3 and the combination therapy exhibited significantly smaller tumor volumes and mases (**Fig. 3G, H, I, J**) compared to control groups. Combination therapy produced enhanced tumor suppression relative to single-agent treatment. Importantly, body weight remained stable throughout the 3-week treatment (**Suppl. Fig. 3G**), and serum analyses revealed no kidney or liver toxicity (**Suppl. Fig. 3H**). Histological evaluation of heart, lung, liver, and colon tissues confirmed the absence of degenerative inflammatory lesions in all therapeutic groups compared to vehicle. Moderate vacuolar degeneration with nuclear pyknosis was present in the proximal tubular epithelium in the kidneys of the combination therapy group but not in single treatment or vehicle groups. Furthermore, the combination of NH-3 with enzalutamide demonstrated enhanced antitumor effects, supporting the potential use of NH-3 in a c combination therapy that would target both the TRβ and AR pathways.

### Thyroid hormone signaling pathway genes have frequent mutations in PCa patients

Genetic alterations in prostate tissue samples from a cohort of treatment-naïve PCa patients who underwent radical prostatectomy (n=102) were analyzed via whole exome sequencing (Ning et al., 2024). The analysis involved the detection of single nucleotide variants (SNVs) and the assessment of their impact on the associated pathways. Mutations were categorized into low, moderate, and high impact (Ensembl Variant Effect Predictor). No genetic changes were detected in TRβ. However, MED12, a key component of the thyroid hormone receptor-associated protein complex, was among the ten most frequently mutated genes (**Fig. 4A**), aligning with literature findings on its role in PCa (Barbieri et al., 2012; Andolfi et al., 2024). Furthermore, a gene set enrichment analysis using KEGG database annotation revealed that “thyroid hormone signaling” ranked ninth among cancer-related and hormonal signaling pathways most affected by mutations (**Fig. 4B**). Notably, mutations were identified in 79 genes within this pathway, with 74.1% of patients harboring single nucleotide variants (SNVs) in these genes (76 out of 102 cases). In addition, “thyroid hormone synthesis” ranked among the most affected pathways, further indicating the importance of TH signaling in PCa pathophysiology (Krashin et al., 2019).

**FIGURE 4:**
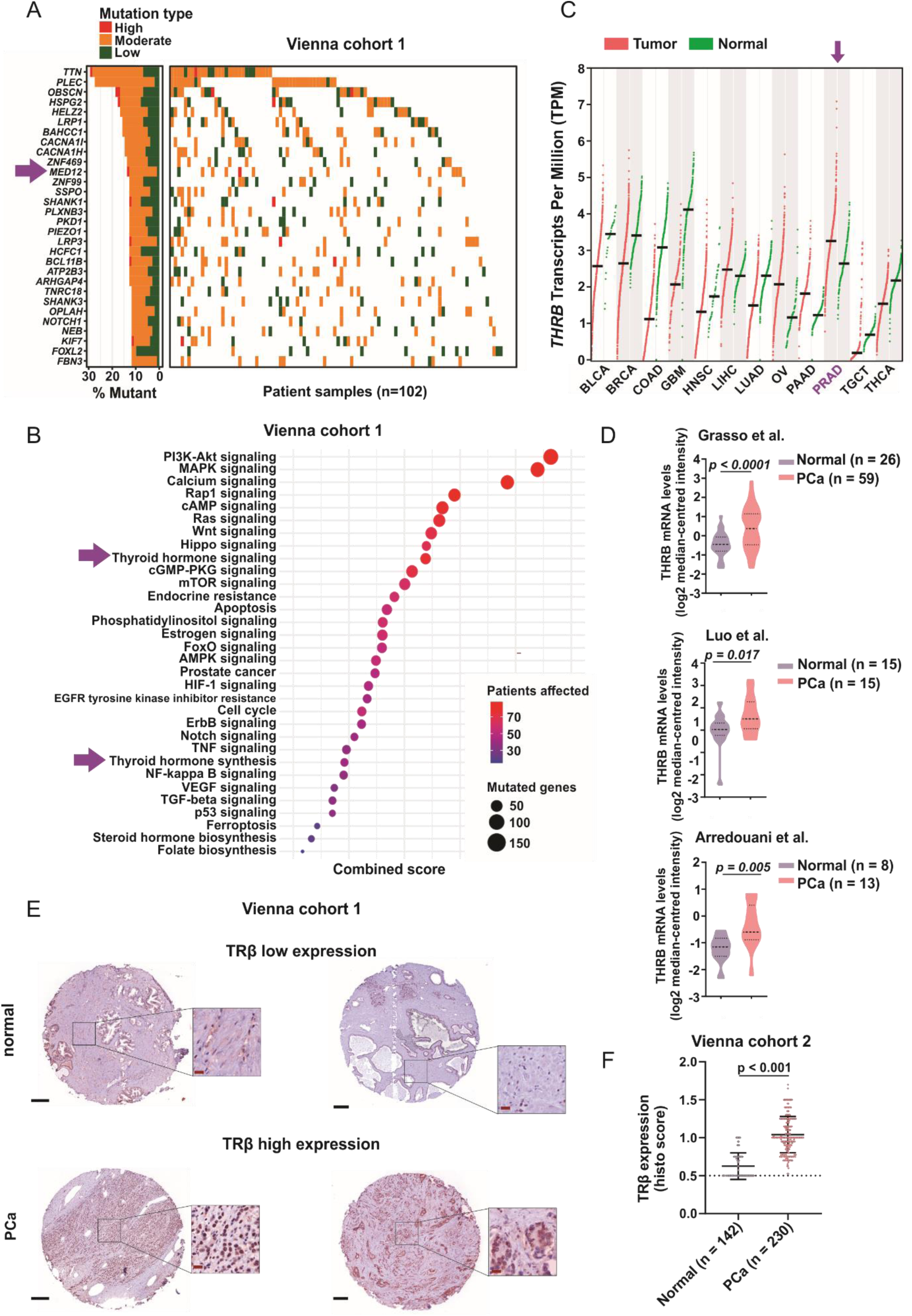
Enrichment of SNVs in thyroid hormone signaling pathway genes and enhanced thyroid hormone receptor beta mRNA and protein expression in PCa patient cohorts. **(A)** Waterfall plot the most frequently mutated genes derived from whole exome sequencing of 102 treatment-naïve patients who had undergone radical prostatectomy. **(B)** Bubble plot of signaling pathways in which the SNV-affected genes from the same PCa patient cohort are enriched. Each bubble represents a pathway; the size corresponds to the number of genes affected by SNVs within that pathway, and the color indicates the number of patients affected. The x-axis represents the combined score, representing the significance and impact of the pathway mutations. Mutation impact ratings (high, moderate, low) were determined according to the Ensembl Variant Effect Predictor (VEP) classification. (C) *THRB* transcript levels across different cancer types (BLCA – Bladder urothelial carcinoma, BRCA – Breast invasive carcinoma, COAD – Colon adenocarcinoma, GBM – Glioblastoma multiforme, HNSC – Head and neck squamous cell carcinoma, LIHC – Liver hepatocellular carcinoma, LUAD – Lung squamous cell carcinoma, OV – Ovarian serous cystadenocarcinoma, PAAD – Pancreatic adenocarcinoma, PRAD – Prostate adenocarcinoma, TGCT – Testicular germ cell tumors, THCA – Thyroid carcinoma), source: GEPIA. Patient numbers in each cohort are listed in the supplementary table. (D) *THRB* mRNA levels extracted using the Oncomine database from three different datasets: Grasso et al. (Normal n = 26, PCa n = 59), Luo et al. (Normal n = 15, PCa n = 15), Arredouani et al. (Normal n = 8, PCa n = 13). **(E)** Representative IHC staining of TRβ in normal prostate and PCa tissue, low expression depicts H-score 20, high expression depicts H-score 300, scale bar 100 µm. **(F)** Histological scoring of IHC staining of TRβ sowing significantly higher TRβ expression in PCa patients (n = 230 slides) compared to healthy control tissue (n = 142 slides).

We also analyzed *THRB* mRNA levels in publicly available human PCa transcriptome datasets. In Gene Expression Profiling Interactive Analysis (GEPIA), *THRB* transcript levels in tumors were higher compared to normal counterparts (**Fig. 4C**). From the Oncomine database, upregulation of *THRB* mRNA was observed across three distinct PCa cohorts compared to normal prostate tissues (**Fig. 4D**). We also examined expression data from primary PCa (n=739) and normal prostate tissues (n=174). We observed that *THRB* transcript levels were again upregulated when comparing primary tumors to controls in Prostate Cancer Transcriptome Atlas (**Suppl. Fig. 4A**) (Bolis et al., 2021).

TRβ protein levels were assessed in an independent cohort of PCa biopsies using immunohistochemical (IHC) staining and scoring (Gleason score scores 3-7) to complement the transcriptomic data. Significantly higher TRβ protein expression was observed in malignant tissues compared to adjacent normal tissues (**Fig. 4F**), corroborating the transcriptomic findings and suggesting a tumor-promoting role for TRβ (Krashin et al., 2019). Taken together, the above findings reveal widespread alterations in the thyroid hormone signaling pathway at the mRNA and protein levels in PCa. Elevated TRβ expression correlates with tumor occurrence, reinforcing its potential role as a biomarker and therapeutic target in PCa.

## DISCUSSION

The identification of the thyroid hormone receptor β (TRβ) as a critical regulator of PCa progression marks a major advance in our understanding of hormonal modulation in cancer biology. Despite its recognized importance, the effects of using specific agonists or antagonists to target TRβ in PCa have not been thoroughly investigated. Here, we have tested the hypothesis that TRβ plays a central role in PCa cell growth and progression. We have shown that specific blockade of TRβ inhibits AR-driven and other signaling pathways that are associated with tumor proliferation. The activity of TRβ plays a role in AR expression, thereby affecting its downstream targets and contributing to the regulation of cellular growth and survival. These findings position TRβ as a novel and promising therapeutic target in PCa.

TRβ has emerged as a versatile and clinically relevant pharmacological target, offering promising therapeutic opportunities through its activation and inhibition. It has dual functionality—either activating or repressing gene expression— which allows precise modulation of signaling pathways involved in diverse pathophysiological conditions, including metabolic diseases and cancer. The selective TRβ agonist resmetirom highlights the therapeutic potential of TRβ activation in hepatocytes. It was effective in treating non-alcoholic steatohepatitis (NASH) in a Phase 3 trial (MAESTRO-NASH) (Harrison et al., 2024). Resmetirom achieved NASH resolution and fibrosis improvement (Guirguis et al., 2024), leading to FDA approval in March 2024 (Keam, 2024). These reports underscore the potential of TRβ activation for metabolic regulation and open avenues for the repurposing of TRβ agonists in diseases in which stimulation of TRβ leads to beneficial changes of gene expression. In contrast to TRβ activation, its inhibition offers a novel strategy for targeting cancers in which TRβ activity contributes to tumor aggressiveness, as in this study through crosstalk with AR-driven PCa growth. This crosstalk and other mechanisms induced by TRβ remain to be elucidated. The selective TRβ antagonist NH-3 competitively binds to the ligand binding domain (LBD) with high affinity and effectively antagonizes T3 activity in luciferase assays (Lim et al., 2002; Shah et al., 2008). Demonstrating specificity and physiological effects characteristic of TRβ blockade, NH-3 has shown broad applicability across human cell lines and animal models (Grover et al., 2007; Ma et al., 2017, Ferrara and Scanlan, 2020; Walter et al., 2021). In this study, NH-3 was rigorously tested for specificity to rule out interference with other nuclear receptors, including AR-related pathways that are critical in PCa (Hantusch et al., 2024). Molecular docking confirmed its selective binding to TRβ. While these findings support the specificity of NH-3, potential off-target effects cannot be completely excluded and warrant further investigation.

We observed that the action of NH-3does not lead to TRβ downregulation at the protein level in PCa cells, thus preserving receptor integrity. This aligns with previous reports that NH-3 disrupts TRβ activation by promoting the release of the corepressors NCOR1 and SMRT without recruiting coactivators such as GRIP-1, TRAP220, or SRC-1 (Nguyen et al., 2002; Lim et al., 2002; Harrus et al., 2018). This mechanism supports a model in which NH-3 disrupts TRβcoactivator interactions and inhibits downstream signaling pathways. We confirmed that NH-3 exhibited potent anti-proliferative activity in both hormone-depleted and hormone-supplemented conditions, demonstrating that its effects are independent of ligand depletion. The delayed onset of action suggests involvement in the regulation of protein synthesis and broader transcriptional control. In PCa models, NH-3 significantly reduced cell proliferation, supporting its role as a promising therapeutic agent for AR-driven and CRPC.

AR has a short half-life and requires prolonged receptor occupancy for stability (Zhou et al., 1995). T3 stabilizes AR independently of androgens, and consequently, we observed AR degradation under hormone-free conditions. in addition, T3 enhances AR signaling and stimulates PSA expression and secretion (Aksoy et al., 2021) and affects AR expression in several models, including Sertoli cells in hypothyroid rats (Panno et al., 1996) and Harderian gland cells in male hamsters (Esposito et al., 2002). In LNCaP cells, T3 increased KLK3 mRNA levels (Esquenet et al., 1995; Zhang et al., 1999; Zhu and Young, 2001), while knockdown of CRYM, a T3 scavenger protein, reduced PSA secretion (Aksoy et al., 2021). PSA, a marker of PCa progression, correlates with sustained AR activity, and its suppression reflects the efficacy of androgen deprivation therapy (ADT). Elevated serum T3 levels in PCa patients with high PSA (Lehrer et al., 2001) and decreased TH levels in ADT-treated patients (Morote et al., 2005) suggest a systemic interplay between TH and AR signaling. *In silico* promoter studies further suggest a combined or mutual regulation of AR- and TR-regulated genes by DHT and T3 (Flood et al., 2013), highlighting the role of T3 in supporting AR-driven processes and emphasizing its relevance in PCa progression. Consistent with this, we showed that NH-3 blockade of TRβ downregulated AR and its targets, Nkx3.1 and PSA, in both androgen-dependent LNCaP and independent PTEN-positive 22Rv1 cells. Synergy maps revealed strong sensitivity of LNCaP cells to low doses of enzalutamide, with peak effects observed at mid-range concentrations when combined with NH-3. In 22Rv1 cells, enzalutamide acted as an enhancer, amplifying ability of NH-3 to downregulate AR signaling. In vitro experiments confirmed that NH-3 effectively inhibits AR activity and reduces Nkx3.1 and PSA expression in PCa cells.

NH-3 treatment led to a significant reduction of PCa tumor growth in NSG xenograft mouse models, consistent with other studies showing that modulation of various NRs has profound effects on tumor dynamics in vivo (Zhang et al., 2024, Xu et al., 2020, Wang et al., 2018). Treatment with NH-3 for three weeks did not cause recognizable side effects, although long-term studies are required to assess potential adverse effects.

Whole exome sequencing (WES) revealed that early stage PCa harbors mutations in previously described genes such as SPOP and MED12 (Barbieri et al., 2012), validating that our cohort is representative. In contrast to previous reports suggesting a tumor-suppressive role of THRB due to deletions and mutations in various cancers (Gonzalezsancho, 2003; Cheng, 2005; Chan and Privalsky, 2009), we did not detect significant SNVs in THRB, suggesting an intact TRβ structure. Nevertheless, 75% of treatment-naïve PCa samples showed perturbations in TH signaling pathways, suggesting an early selective advantage conferred by TH activity. To date, alterations in thyroid hormone signaling have rarely been described in PCa, as most transcriptomic and proteomic analyses assess expression rather than function.

Our analyses revealed that *THRB* mRNA and TRβ protein levels were upregulated in tumor samples compared to normal controls in several cohorts. Altered TRβ expression has also been observed in other cancers, including colorectal cancer (Hörkkö et al., 2006), breast cancer (Heublein et al., 2015; Shao et al., 2020) and head and neck squamous cell carcinoma (Schnoell et al., 2021). These studies also reported an adverse survival correlation with nuclear or cytosolic TRβ localization, underscoring the need to distinguish between nuclear and cytosolic TRβ expression and highlighting the role of intracellular T3 availability in shaping the regulatory effect of TRβ on tumor progression.

In this study, we demonstrate for the first time that activated TRβ significantly affects AR activity and promotes PCa cell growth. Specifically, we show that TRβ activation drives PCa cell growth and regulates AR-dependent processes. Importantly, we highlight the therapeutic potential of targeting TRβ with the compound NH-3, which shows high efficacy against PCa cells. Our findings underscore the broader significance of TRβ activation as a fundamental oncogenic mechanism, particularly in the context of CRPC. The synergistic inhibition of cell proliferation observed with NH-3 and enzalutamide in both LNCaP and 22Rv1 cells suggests a promising combinatorial approach to enhance AR inhibition. Notably, NH-3 demonstrated potent activity under ADT conditions, where impaired AR signaling amplifies the dependence on TRβ-mediated pathways, positioning TRβ as an essential factor for sustained cancer progression.

We propose in this study that TRβ-selective antagonists, such as NH-3, alone or in combination with enzalutamide, represent a novel and effective therapeutic strategy for PCa, particularly CRPC. The dual targeting of AR through distinct mechanisms may reduce the likelihood of resistance development, a pervasive challenge in current treatment paradigms. Nevertheless, future studies are needed to validate these findings and elucidate the full spectrum of TRβ oncogenic roles and its potential as a therapeutic target. This work fundamentally advances our understanding of PCa biology by highlighting TRβ as a critical factor in cancer progression, offering new avenues for combination therapies to address unmet clinical needs in advanced PCa.

## MATERIALS AND METHODS

### PCa patient cohorts and characteristics

#### Vienna cohort 1

This study utilized data previously published by Ning J. et all (DOI: 10.7150/thno.96921). The waterfall plot was generated in R using the GenVisR package (Skidmore et al., 2016)." Skidmore ZL, Wagner AH, Lesurf R, Campbell KM, Kunisaki J, Griffith OL, Griffith M (2016). " GenVisR: Genomic Visualizations in R." Bioinformatics, 32, 3012-3014.

#### Vienna cohort 2

Formalin-fixed, paraffin-embedded (FFPE) samples were collected from patients who had radical prostatectomy at the Department of Pathology, Medical University of Vienna, Austria, and the Institute of Pathology, Tuebingen, Germany. Immunohistochemistry (IHC) was performed following standard protocols, and protein expression was quantified as previously described (doi: 10.1371/journal.pone.0100822). The following TRβ antibody was used: Cat# 209-301-A96, 1:100, Rockland. Tissue microarrays (TMAs) were evaluated by four certified pathologists according to a 4-point scale for staining intensity (0-3) and a 4-point scale for the number of stained cells (0-3).

### Publicly available databases

*THRB* mRNA expression levels were obtained from The Cancer Genome Atlas (TCGA) from the following datasets: prostate adenocarcinoma (PRAD), bladder urothelial carcinoma (BLCA), breast cancer (BRCA), colon adenocarcinoma (COAD), glioblastoma multiforme (GBM), head and neck squamous cell carcinoma (HNSC), liver hepatocellular carcinoma (LIHC), lung adenocarcinoma (LUAD), ovarian carcinoma (OV), pancreatic adenocarcinoma (PAAD), testicular germ cell tumors (TGCT), thyroid carcinoma (THCA). For tumor compared to normal sample analysis, the GEPIA tool was used (http://gepia.cancer-pku.cn). The ENSG00000151090 identifier was used for the *THRB* gene (Ensembl version 109). Expression levels are reported as normalized Transcripts Per Million (TPM) values, which were derived from RNA-sequencing experiments conducted from tissue samples.

Gene expression data for *THRB* mRNA were extracted from various prostate cancer datasets (Grasso Prostate, Luo Prostate 2, Arredouani Prostate), including normal and tumor samples, using in the Oncomine Research Premium Edition database (Thermo Fisher, Ann Arbor, MI) (10.1016/s1476-5586(04)80047-2).

### Principal component analysis (PCA)

To visualize the expression levels of *THRB* in PCa samples from individual patients and healthy tissue, we examined the data set from Bolis et al., 2021, and used the autoplot function of the R package ggplot2. Boxplot representations of *THRB* expression in PCa and healthy normal tissue of stated samples show the median (center line), the upper and lower quartiles (the box), and the range of the data (the whiskers), including outliers. Significance was determined by an unpaired, two-tailed t-test using the R package ggplot2.

### Cell culture

#### Cell lines

Human RWPE1, BPH1, LNCaP, 22Rv1, DU145 and PC-3, prostate cancer cell lines were obtained from the American Type Culture Collection (ATCC, Manassas, VA, USA) and cultured in RPMI 1640 medium supplemented with 10% fetal bovine serum (FBS) and 1% penicillin-streptomycin (Gibco, #11875093, #15140122 and #26140079, respectively). The cells were maintained in a humidified incubator at 37°C with 5% CO_2_. All cell lines were tested to be free of mycoplasma infection.

### Resazurin assay

Cells were seeded in 96-well plates at a density of 2-3.5×10 3 cells per well. Cell were cultured in full media (FM: RPMI, 10% FCS, 1% PenStrep), hormone-free media (HFM: phenol red-free RPMI (Gibco, #11835030), 10% charcoal-stripped FBS (Gibco, #12676029), 1% PenStrep) and hormone-free media with DMSO (HFM+DMSO). 10 µM bortezomib (MedChemExpress, HY-10227) was as positive control for cell death. After respective treatment time points, Resazurin reagent (Santa Cruz, # sc-206037) was added to each well and incubated for 3 h at 37°C. Light emission of the reduced fluorescent product resorufin was measured using a TECAN Synergy H1 plate reader (Tecan Group) at 570 nm. The fluorescence intensity was recorded, and the relative cell viability or proliferation was calculated by comparing the fluorescence signals of the treated wells to those of the control wells. Statistical analysis was performed using one-way ANOVA and significance was defined at p < 0.05.

### Drug treatments

Selective TRβ inhibitor NH-3 (synthesized in the laboratory of Patrick Gunning, University of Toronto), 1-850 (Sigma-Aldrich, 609315), enzalutamide (Axon, MDV 3100), bicalutamide (MedChemExpress, HY-14249; Sigma-Aldrich, B9061), and T3 (MedChemExpress, HY-A0070A) were diluted in DMSO and treatments were performed in FM or HFM as described above. Each treatment condition was performed in triplicates.

### IncuCyte cell proliferation assay

The IncuCyte® S3 Live-Cell Analysis System (Sartorius, Michigan, MI, USA) was utilized for real-time kinetic monitoring of cell growth upon treatments as described previously according to the manufacturer’s protocol. Phase-contrast images were automatically acquired from multiple fields per well every 2 hours for the duration of the experiment. Cell proliferation was quantified using the IncuCyte integrated analysis software. The software’s built-in algorithm was used to calculate cell monolayer density based on the phase-contrast images. This metric provides a measure of cell confluence and growth over time.

### Ki67 analysis

Ki-67 staining was performed to analyze the proliferative activity of target cells. The cells were fixed with 70% ethanol and incubated at −20°C for 1 hour. After two washes with Cell Staining Buffer (#420201, BioLegend®), cells were resuspended and incubated with a Ki-67 antibody (#350501, BioLegend ®) for 30 min at room temperature in the dark. Following two additional washes, cells were incubated with a secondary antibody, goat anti-mouse IgG (H+L) Cross-Adsorbed Secondary Antibody, Alexa Fluor™ 488 (# A-11001, Invitrogen), for 20-30 minutes in the dark. Stained cells were resuspended in cell staining buffer for flow cytometric analysis. All samples were acquired with a BD FACSCanto II (Becton Dickinson, Franklin Lakes, NJ, USA) and analyzed with FlowJo 7.6 software (BD Biosciences).

### Competitive radioligand binding assay

Radioligand binding assays for TRβ (#285940), RXRα (#269500), RXRβ (#269540), AR (#933), ERα (#5484), ERβ (#226050), GR (#469), and PR (#2341) was performed at Eurofins (https://www.eurofins.com/) according to the company standards. To evaluate the capacity of NH-3 (MUV-1, PT# 1281206) to interfere with radiolabeled ligand binding, increasing concentrations of NH-3 were applied, and radioactive signals were monitored. IC50 values were determined by a non-linear, least squares regression analysis using MathIQTM (ID Business Solutions Ltd., UK). Inhibition constants (Ki) values were calculated using the equation of Cheng and Prusoff (Cheng, Y., Prusoff, W.H., Biochem. Pharmacol. 22:3099-3108, 1973) using the observed IC50 of the tested compound, the concentration of radioligand employed in the assay, and the historical values for the KD of the ligand (obtained experimentally at Eurofins Panlabs, Inc.). Hill coefficient (nH), defining the slope of the competitive binding curve, was calculated using MathIQTM.

### *In vitro* [^68^Ga]PSMA uptake

LNCaP cells were treated NH-3 as described above, washed with PBS, trypsinized and centrifuged. 100 000 cells were incubated on 96-well filter plates (MADVN6550, Merck Millipore, Darmstadt, Germany) with 36 kBq [68Ga]PSMA-11 for 1 hour in an incubator (humidified atmosphere, 37°C, 5% CO2). The radiotracer was freshly produced on-site before the experiment. To assess unspecific binding, the filter plates were incubated with the radiotracer alone. After incubation, the cells were washed by vacuum filtration with PBS (2 × 200 µL) through the plate. The filters were transferred into tubes using a commercial punch kit and measured in a gamma counter (Wizard 2, PerkinElmer, Waltham, MA, USA). Radiotracer uptake was quantified as the percentage of added radioactivity per 100 000 cells. Unspecific binding in all samples was <0.1 %.

### Synergy analysis

Synergy was calculated based on resazurin assay as described above. The degree of synergy was quantified using the Loewe Additivity model. The Loewe model assumes that the expected effect of a drug combination is as if a drug is combined with itself, considering the full dose-response relationships of the individual drugs. The four-parameter logistic regression (LL4) was used as the curve-fitting algorithm. This model provides a more accurate and nuanced interpretation of drug interactions, incorporating parameters such as EC50 and dose-response curves. A Loewe synergy score is derived to evaluate the combined effects, where deviations from the expected additivity indicate the presence of synergistic or antagonistic interactions.

### Protein isolation and Western blot analysis

Cells were harvested by trypsinization and pelleted by centrifugation at 500 x g for 5 min. Cell pellets were washed with ice-cold PBS and lysed in RIPA buffer (Sigma-Aldrich, St. Louis, MO, USA, #D8418) supplemented with protease (cOmplete™ Mini Proteasehemmer-Cocktail, Roche, #11836153001) and phosphatase inhibitors (PhosSTOP™, Roche, # 4906845001). The cell lysates were incubated on ice for 15 min. Subsequently, the lysates were centrifuged at 20 000 g for 20 min at 4°C. The supernatants were collected, protein concentrations were determined by BCA (Thermo Scientific™, 23227) method and stored at −80°C. Equal amounts of protein samples were separated by SDS-PAGE using 10% polyacrylamide gels. Proteins were then transferred onto nitrocellulose membranes (Cytiva, #10600001) using a Trans-Blot Turbo Transfer System (Bio-Rad, Hercules, CA, USA). The membranes were blocked with 5% BSA and incubated with primary antibodies (table below) overnight at 4°C. After 3×5 min washing, the membranes were incubated with HRP-conjugated secondary antibodies and visualized using an ECL detection system (Clarity™ Western ECL Substrate, #1705060) using the ChemiDoc XRS+ (Bio-Rad) system.

| Antibody | Dilution | Cat. number and company |
| --- | --- | --- |
| TRβ | 1:1000 in 5% BSA | Abx001332, Abbexa |
| AR | 1:1000 in 5% BSA | ab74272, Abcam |
| PSA | 1:1000 in 5% BSA | 5365s, Cell Signaling |
| Nkx3.1 | 1:1000 in 5% BSA | #83700, Cell Signaling |
| Tubulin | 1:5000 in 5% BSA | #86298, Cell Signaling |
| Actin | 1:5000 in 5% BSA | #3700, Cell Signaling |

### Mouse xenografts

Animal work was carried out according to ethical standards and was approved by the Medical University of Vienna and by the Austrian Federal Ministry of Science, Research, and Economy (GZ66.009/377-V/3b/2018). NSG mice were provided by the Institute of in vivo and in vitro Models (University of Veterinary Medicine, Vienna) and by the Core facility Laboratory Animal Breeding and Husbandry (Medical University, Himberg). For 22Rvi xenografts, NOD.Cg-Prkdcscid Il2rgtm1Wjl Tg(CMV-IL3,CSF2,KITLG)1Eav/MloySzJ mice were used, for LNCaP xenografts NOD SCID mice were used. To establish xenografts, NSG mice were injected subcutaneously with 22Rv1 (2 Mio. cells per flank) and LNCaP (4 Mio. cells per flank). Tumor growth was monitored biweekly by measuring tumor dimensions with a caliper. Once tumors reached a palpable size, mice were randomly assigned to treatment groups. Vehicle formulation was as follows: (5% (v/v) ethanol, 5% (v/v) Kolliphor EL, 30% (v/v) propylene glycol and 20% (w/v) HP-beta-CD in PBS (pH 7.4). Mice were treated daily with NH-3, enzalutamide, or both, diluted in vehicle and administered intraperitoneally (i.p.) or by oral gavage (o.g.). Control mice received vehicle only. During the treatment period, mice were closely monitored for any signs of distress or adverse side effects. Tumor growth was monitored, and changes in tumor volume were recorded by measuring the length (L) and width (W) using a caliper. Tumor volume was calculated using the formula V = (L × W^2) / 2. At the end of treatment, mice were euthanized, blood samples were taken by heart puncture, and xenograft tumors and organs were excised for further analysis. Statistical analysis was performed, and significance was determined at a *P-*value < 0.05.

### Histological analysis of tumor xenografts

Organs were fixed overnight in 4% phosphate-buffered formaldehyde solution (Roti® Histofix, Carl Roth), dehydrated, paraffin-embedded, and cut into 2 µm consecutive organ or tumor sections. Sections were subsequently stained with Hematoxylin & Eosin G (according to standard routine protocol), Ki-67 (Cell Marque, SP6), and THRβ (Rockland, 209-301-A96). Slides were scanned using Vectra Polaris™ (Akoya Bioscences). Organ toxicity evaluation was performed by an animal pathologist. IHC results were consulted with two independent expert pathologists.

### Blood parameters evaluation

Mouse blood collected by heart puncture was added to EDTA-tubes (Mini-Collect K3EDTA tubes). Plasma was obtained by centrifuging the whole blood for 20 min at 1000 x g. Blood parameters were assessed with an animal blood counter (Vet abc, scil animal care; Hitachi/Roche Cobas 4000 c311 Analyzer). Plasma concentrations of blood urea nitrogen (BUN), aspartate aminotransferase (AST) and alanine aminotransferase (ALT) were determined using a laboratory chemistry analyzer (IDEXX VetTest 8008, IDEXX Laboratories) and specific assays (04467388190, 04467493190, 04460715 190, cobas).

### Computational modeling

Protonation states of the amino acid side chains were estimated by using the H++ program (Anandakrishnan et al., 2012). The protein was solvated by water molecules, and a number of Na+ and Cl-ions that represent 0.154 M NaCl solution, and further sodium ions to neutralize the charge of the protein. To describe the intra- and intermolecular interactions within the periodic simulation box, the charmm36 force field was used for the biomolecules, the ligands, and the ions (Best et al., 2012), whereas for water the TIP3P model was employed (Jorgensen et al., 1983). Bonds to hydrogen atoms were constrained throughout the simulations. The ligand topologies were generated by the CGenFF web server was used (Vanommeslaeghe et al., 2010). Molecular dynamics simulations were performed with Gromacs 2024.3 (Abraham et al., 2024; Abraham et al., 2015; Pronk et al., 2013; Berendsen et al., 1995). To control the temperature and the pressure in the simulations, a modified Berendsen thermostat (reference temperature 310 K, time constant 0.1 ps) and a Berendsen barostat (reference pressure 1 bar, time constant 2 ps) were applied. The timestep was chosen to be 2 fs. Molecular docking calculations were performed at the Swissdock webserver (Bugnon et al., 2024; Grosdidier et al., 2011; Röhrig et al., 2023; Zoete et al., 2016) using Autodock Vina (Eberhardt et al., 2021; Trott and Olson, 2010).

### Statistical analyses

For datasets with normal distribution, the Student’s *t*-test was used for group comparison. If the distribution was not normal, the Mann-Whitney *U* test was used. Statistical significance was defined as p < 0.05 unless stated otherwise. Significance levels are indicated as follows: *p < 0.05, **p < 0.01, ***p < 0.001, ****p < 0.0001.

## Author contributions

Conceptualization and design: A.F., B.H., L.K.

Resources: L.K., E.D.A, G.F., T.W., G.T., J.S.Y., Y.S.R., O.O.O., P.T.G., M.D., J.N.

Acquisition of data: A.F., D.P., M.S., C.P., A.W., T.W., O.A.

Methodology: D.P., M.S., A.M.M., M.V., B.S. O.H., V.B., G.W.

Project administration: A.F., B.H., L.K.

Formal analysis: A.F., S.H., C.P., H.N., T.R., M.S., C.S.

Interpretation of data: A.F., B.H., L.K.

Study supervision: B.H., L.K.

Funding acquisition: L.K.

Scientific advice: M.S., R.K., G.B., J.P., D.H., O.M.

Figure preparation: A.F.

Writing - original draft: A.F., B.H., L.K.

All authors read and approved the final manuscript.

## ACKNOWLEDGMENTS

L.K. acknowledges the support from MicroONE, a COMET Modul under the lead of CBmed GmbH, which is funded by the federal ministries BMK and BMDW, the provinces of Styria and Vienna, and managed by the Austrian Research Promotion Agency (FFG) within the COMET—Competence Centers for Excellent Technologies—program. Financial support was also received from the Austrian Federal Ministry of Science, Research and Economy, the National Foundation for Research, Technology and Development, and the Christian Doppler Research Association, as well as Siemens Healthineers for their financial and scientific support. L.K. was also supported by a European Union Horizon 2020 Marie Sklodowska-Curie Doctoral Network grants (FANTOM, n. P101072735 and eRaDicate, n. 101119427) the Christian-Doppler Lab for Applied Metabolomics (CDL-AM), and the Austrian Science Fund (grants FWF: P26011, P29251, P 34781 as well as the International PhD Program in Translational Oncology IPPTO 59.doc.funds). Additionally, this research was funded by the Vienna Science and Technology Fund (WWTF), grant number LS19-018. L.K. is a member of the European Research Initiative for ALK-Related Malignancies (www.erialcl.net).

The financial support for O.H. by the National Research, Development and Innovation Office through the project OTKA-FK 138823 is gratefully acknowledged. Furthermore, O.H. is grateful for the support from the János Bolyai Research Scholarship of the Hungarian Academy of Sciences, and the ÚNKP-22-5 and ÚNKP-23-5 New National Excellence Program from the National Research, Development and Innovation Fund. The authors thank the Vienna Supercomputing Center and the Governmental Information Technology Development Agency (KIFÜ) for the CPU time that has been used for this project.

We acknowledge our collaborators: Enikö Kallay (Medical University of Vienna), Christopher Roberts and Yi Zhao (University of Nottingham). Medical University of Vienna Core Facility Imaging I acknowledged for providing utilization of Vectra Polaris. We also acknowledge the use of BioRedner.com for creating Fig. 1F, 2A, 2F, 3F and Suppl. Fig. 1C.

