## Supplementary figures for "Thyroid Hormone Receptor Beta Signaling is a Targetable Driver of Prostate Cancer Growth"

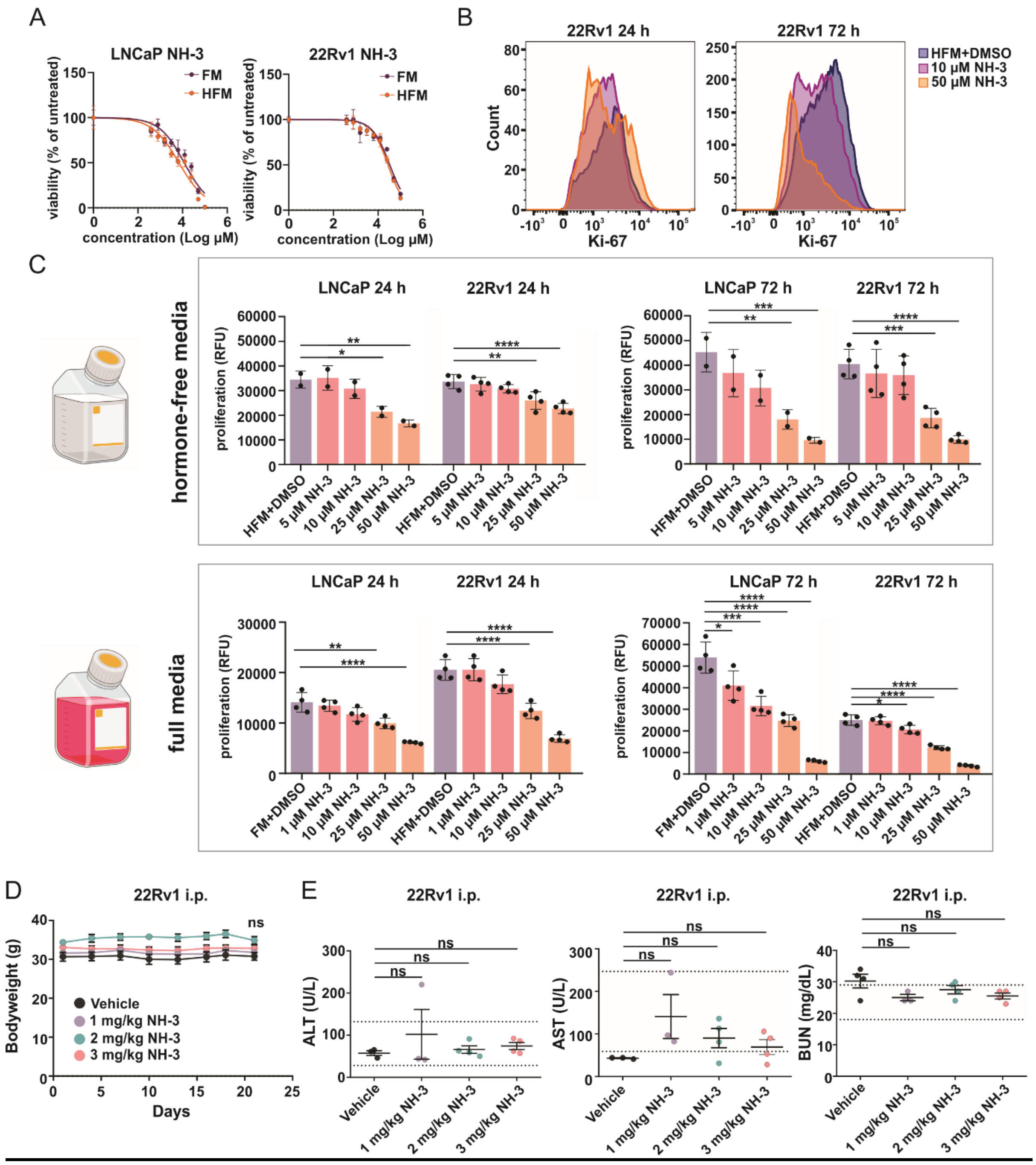


**Supplementary Figure 1. (A)** Dose-response curves of NH-3 treated LNCaP and 22Rv1 cells after 72h treatment. **(B)** Reduced proliferation of 22Rv1 cells induced by increasing NH-3 treatment, as seen by a shift towards Ki-67-low expressing cells. **(C**, upper panel**)** Reduced proliferation of NH-3 treated LNCaP and 22Rv1 in Resazurin assay in hormone-free conditions and **(C**, bottom panel) in full media. Created with BioRender.com. **(D)** No alterations of body weight in 22Rv1 xenograft mice upon 1, 2, 3 mg/kg NH-3 treatment compared to controls. **(E)** Unchanged liver and kidney parameters in 22Rv1 xenograft mice upon 1, 2, 3 mg/kg NH-3 treatment compared to controls. Mean ± SD, *p < 0.05, **p < 0.01, and ***p < 0.001. ALT - alanine aminotransferase, AST - aspartate aminotransferase, BUN - blood urea nitrogen.

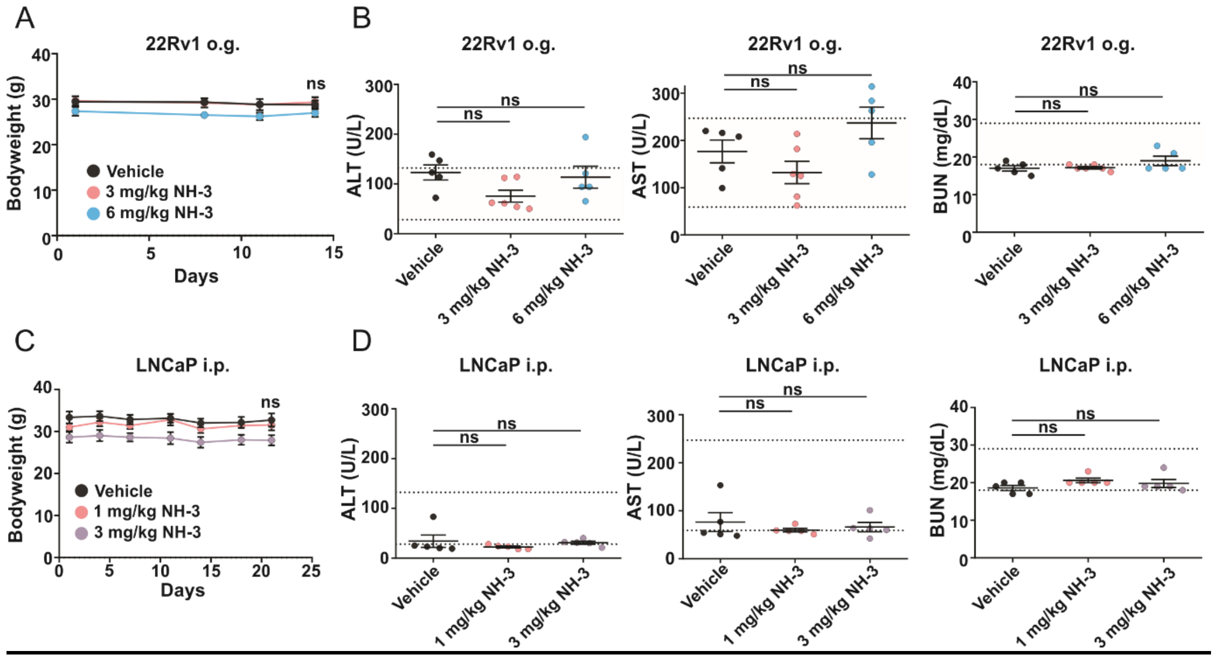


**Supplementary Figure 2.**

**(A)** Body weight of 22Rv1 xenograft mice treated with vehicle, 3 and 6 mg NH-3/kg/day by o.g. administration **(B)** Liver and kidney parameters in sera of 22Rv1 xenograft mice treated with 3 and 6 mg NH-3/kg/day by o.g. administration **(C)** Body weight of LNCaP xenograft mice treated with vehicle, 1 and 3 mg NH-3/kg/day by i.p. administration. **(D)** Liver and kidney parameters in sera of LNCaP xenograft mice treated with vehicle, 1 and 3 mg NH-3/kg/day by o.g. Mean ± SD, *p < 0.05, **p < 0.01, and ***p < 0.001. ALT - alanine aminotransferase, AST - aspartate aminotransferase, BUN - blood urea nitrogen.

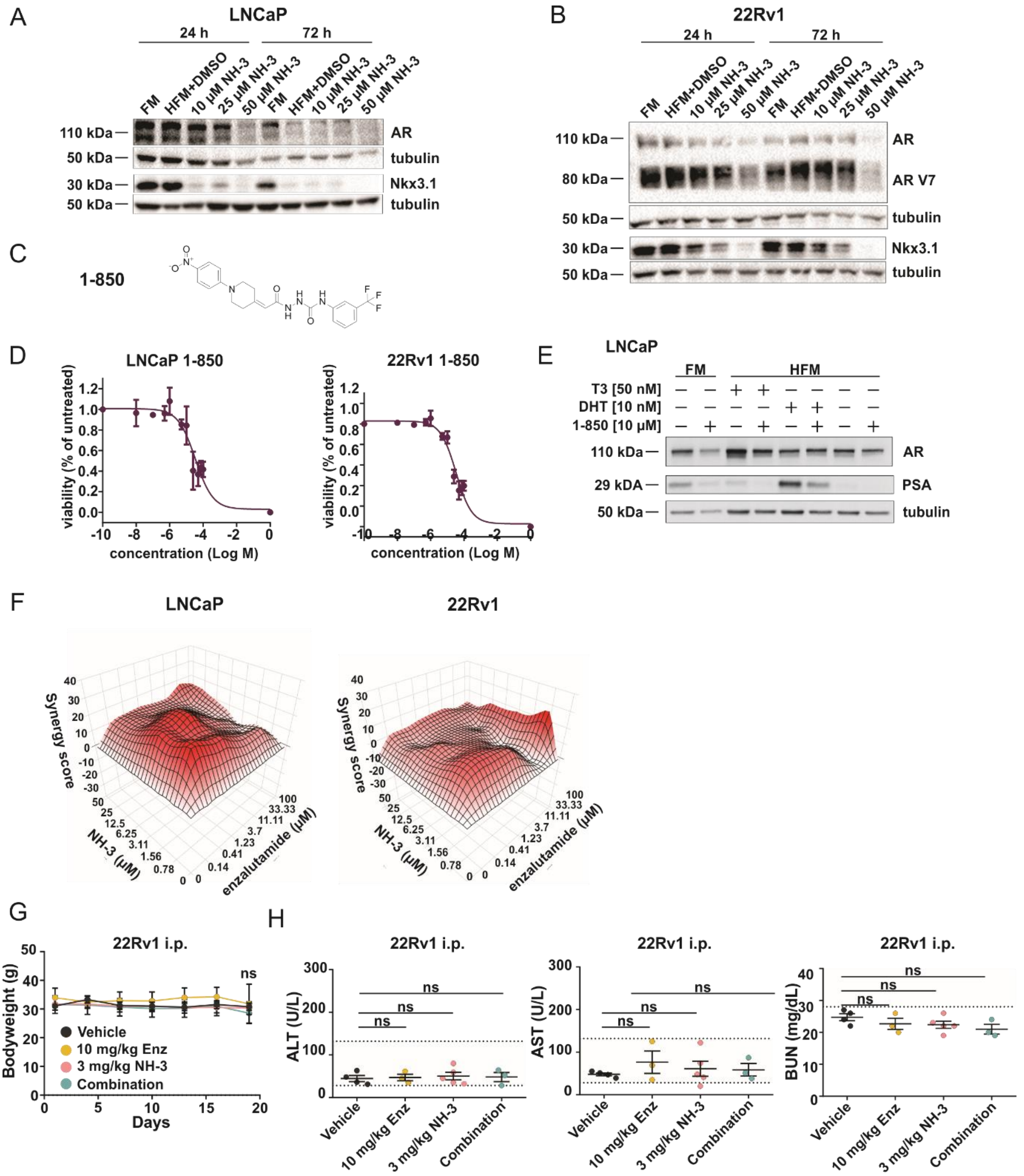


**Suppl. Fig. 3. (A)** Western blot analysis of LNCaP cells treated with increasing NH-3 concentrations after 24 h. Representative blot showing expression of TRβ, AR, PSA, Nkx3.1; GAPDH and tubulin as loading controls. **(B)** Western blot analysis of 22Rv1 cells treated with increasing NH-3 concentrations after 24. Representative protein expression of TRβ, AR, PSA, Nkx3.1; GAPDH and tubulin as loading controls. **(C)** Molecular structure of 1-850. **(D)** IC-50 values for 1-850 in different LNCaP and 22Rv1 cell lines. **(E)** Western blot analysis of AR and PSA protein expression analyzed in LNCaP cells treated with 10 µM 1-850 in FM, HFM or HFM supplemented with 1 nM T3 after 72 hours. β-tubulin was used as loading control. **(F)** Synergy maps of NH-3, enzalutamide and combined treatment, created with "Synergy Finder". **(G)** Body weights of 22Rv1 xenograft mice upon 3 mg/kg/day NH-3, 10 mg/kg/day enzalutamide and combinatory i.p. treatment **(H)** Liver and kidney parameters in 22Rv1 xenograft mice upon i.p. 3 mg/kg/day NH-3, 10 mg/kg/day enzalutamide and combinatory treatment. Mean ± SD, *$p < 0.05$, **$p < 0.01$, and ***$p < 0.001$. ALT - alanine aminotransferase, AST - aspartate aminotransferase, BUN - blood urea nitrogen.

A

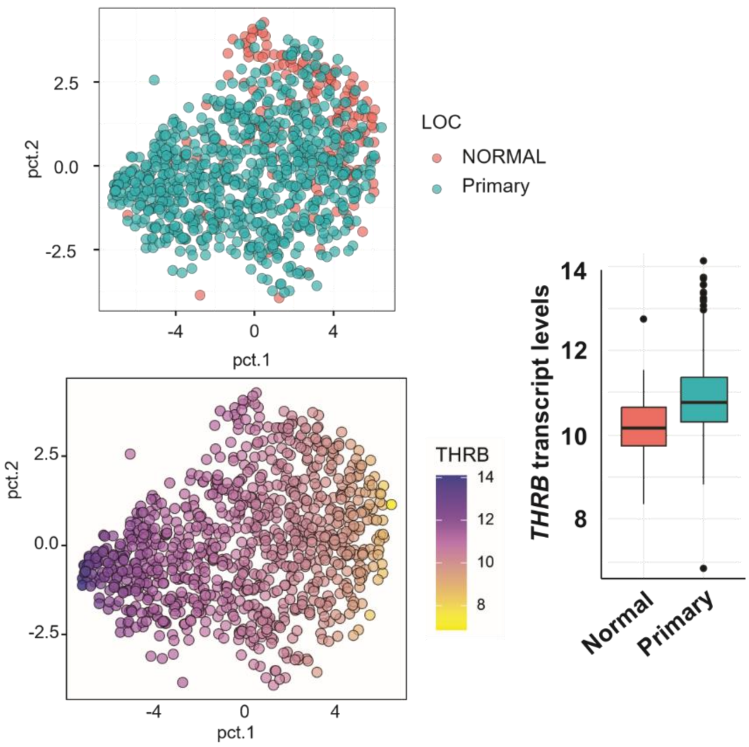


**Suppl. Fig. 4. (A)** PCA representation of expression levels of THRB in subsets of primary PCa (Primary, n=739) and normal prostate tissue (NORMAL, n=174), left panels and box plot representation, right panel.
